# HyDRA: a pipeline for integrating long- and short-read RNAseq data for custom transcriptome assembly

**DOI:** 10.1101/2024.06.24.600544

**Authors:** Isabela Almeida, Xue Lu, Stacey L. Edwards, Juliet D. French, Mainá Bitar

**Author notes:** These authors contributed equally (Juliet D. French, Mainá Bitar). Correspondence: Mainá Bitar, QIMR Berghofer Medical Research Institute, 300 Herston Road, Herston, Australia 4006.

## Abstract

**Background:** Short-read RNA sequencing (RNAseq) has widely been used to sequence RNA from a wide range of different tissues, developmental stages and species. However, the technology is limited by inherent biases and its inability to capture full-length transcripts. Long-read RNAseq overcomes these issues by providing reads that can span multiple exons, resolve complex repetitive regions and the capability to cover entire transcripts. Unfortunately, this technology is still prone to higher error rates. Noncoding RNA transcripts are highly specific to different cell types and tissues and remain underrepresented in current reference annotations. This problem is exacerbated by the dismissal of sequenced reads that align to genomic regions that do not contain annotated transcripts, resulting in approximately half of the expressed transcripts being overlooked in transcriptional studies.

**Results:** We have developed a pipeline, named HyDRA (<u>Hy</u>brid *<u>d</u>e novo* <u>R</u>NA <u>a</u>ssembly), which combines the precision of short reads with the structural resolution of long reads, enhancing the accuracy and reliability of custom transcriptome assemblies. Deep, short- and long-read RNAseq data derived from ovarian and fallopian tube samples were used to develop, validate and assess the efficacy of HyDRA. We identified more than 50,000 high-confidence long noncoding RNAs, most of which have not been previously detected using traditional methods.

**Conclusions:** HyDRA’s assembly performed more than 40% better than a similar assembly obtained with the top-ranked stand-alone *de novo* transcriptome short-read-only assembly tool and over 30% better than one obtained with the best-in-class multistep short-read-only approach. Although long-read sequencing is rapidly advancing, the vast availability of short-read RNAseq data will ensure that hybrid approaches like the one implemented in HyDRA continue to be relevant, allowing the discovery of high-confidence transcripts within specific cell types and tissues. As the practice of performing hybrid *de novo* transcriptome assemblies becomes commonplace, HyDRA will advance the annotation of coding and noncoding transcripts and expand our knowledge of the noncoding genome.

## BACKGROUND

Short-read RNA sequencing (RNAseq) has revolutionized the transcriptomic era due to its high-throughput, affordability and low error rates^1^. However, a limitation of short-read RNAseq lies in its dependency on fragmenting the original transcript molecules. Reassembling and quantifying these sequenced reads, typically of ~50 to ~500 nt in length, still poses significant computational challenges^2^. The majority of RNAseq studies measure the expression of genes and transcripts by mapping the sequenced reads to a reference annotated transcriptome and removing reads that fail to map. Notably, reference transcriptomes such as those annotated by ENSEMBL and GENCODE are far from complete^3^, leaving a large proportion of transcripts unquantified by standard RNAseq analysis methods^3^. To overcome these limitations, a *de novo* custom transcriptome assembly can be performed, to reconstruct the sequenced fragments of transcripts expressed in the sample of interest in substitution of a reference. Mapping sequenced reads onto a custom transcriptome allows the quantitation of expression levels from both annotated and unannotated transcripts^4^. However, to date, only a small fraction of publications make use of *de novo* custom assemblies, accounting for ~1% of the total PubMed publications that use RNAseq. In addition, *de novo* assemblies obtained using only short reads cannot accurately resolve all RNA transcripts^5^.

Long-read sequencing platforms such as those from Pacific Biosciences (PacBio) and Oxford Nanopore Technology (ONT) have the potential to produce full-length transcripts^6^. These platforms can perform end-to-end sequencing of single complementary DNA (cDNA) or RNA molecules, generating long reads that ameliorate the issues caused by transcript fragmentation^7^. ONT platforms include a range of devices in which single molecules thread through a nanopore containing a nanoscale sensor able to detect each nucleotide within a single run^8^. The produced long reads are one order of magnitude longer than typical short reads, providing better resolution of splice junctions, increasing correct isoform identification and the discovery of unannotated transcripts^8^. However, compared to short reads, long reads have much higher base-calling error rates^7,9,10^ and remain a costly and comparatively less used technology. Importantly, although both long-read sequencing and basecaller technologies are continually evolving, most facilities are not equipped to support long-term storage of the large raw data files due to associated costs. Therefore, most users cannot re-call previously sequenced reads when an improved basecalling algorithm is released to increase the quality of long-read data.

Emerging studies have shown that hybrid transcriptome assembly approaches, which integrate short- and long-read RNAseq data, are more accurate than approaches that use data from either method independently^11,12,13,14,15,16^. Considering that the average human transcript length is one kilobase (kb)^17^ and that long-read sequences are on average 1-3 kb in length^18,19^, long reads should capture the majority of human transcripts within a single read and ideally bypass the need for reassembly. However, considering the low quality of long reads^7,9,10^, it is still recommended to correct for intrinsic errors^9^. Although there are different strategies to achieve this, a pre-assembly hybrid error correction using both short and long reads was recently shown to be the best-performing method^10^. Additionally, information from both long and short reads may be integrated at the assembly stage, to help reconstruct different isoforms^20^. To the best of our knowledge, none of the available tools fully benefit from the two types of read integration, but instead adopt either a hybrid-correction-only or a hybrid-assembly-only approach.

Notably, for long noncoding RNAs (lncRNAs), which constitute the largest class of underrepresented RNA transcripts, their lower abundance in bulk tissues and high content of repetitive elements means the assembly challenge is even more pronounced^21,22,23^. The few lncRNA-focused hybrid assembly studies that have been performed indicate that RNAseq data integration can enhance the accuracy and reliability of lncRNA discovery^24,25^. However, no automated method for hybrid *de novo* assembly to date allows for accurate lncRNA discovery.

To address the need for comprehensive discovery of unannotated transcripts, we developed HyDRA (<u>Hy</u>brid *<u>d</u>e novo* <u>R</u>NA <u>a</u>ssembly), a true-hybrid pipeline that integrates short- and long-read RNAseq data for *de novo* transcriptome assembly, with additional steps for lncRNA discovery. Our pipeline combines read treatment, assembly, filtering and parallel quality control (QC) steps to ensure the reconstruction of high-quality transcripts. Comprehensive tests showed that HyDRA outperforms the current best-in-class short-read-only approach^4^. In contrast with long-read sequencing, a vast amount of short-read RNAseq data is readily available for many species, tissues and conditions. Pipelines like HyDRA can make best use of available data in its totality, allowing users to achieve high-quality transcriptome assemblies while long-read sequencing technologies continue to advance. We anticipate that HyDRA will facilitate the generation of tissue-specific custom transcriptomes, providing a valuable resource for expression analyses across different cell types and tissues.

## RESULTS AND DISCUSSION

### Overview of the HyDRA pipeline

We developed HyDRA (Figure 1A), a hybrid pipeline that integrates bulk short- and long-read RNAseq data for generating custom transcriptomes. This is achieved through (i) read treatment steps to correct sequencing errors by treating low-frequency *k*-mers and removing contaminants (e.g. adaptors and reads from ribosomal RNAs), (ii) steps to *de novo* assemble the filtered and corrected reads and further process the resulting assembly, and (iii) optional steps to discover a high-confidence set of lncRNAs supported by multiple machine-learning model predictions (Figure 1B-D). This section and Additional file 1 contain a detailed explanation of HyDRA, including the tools and algorithms underlying each step (Table 1; Additional File 2: Table S1).

**Figure 1.**
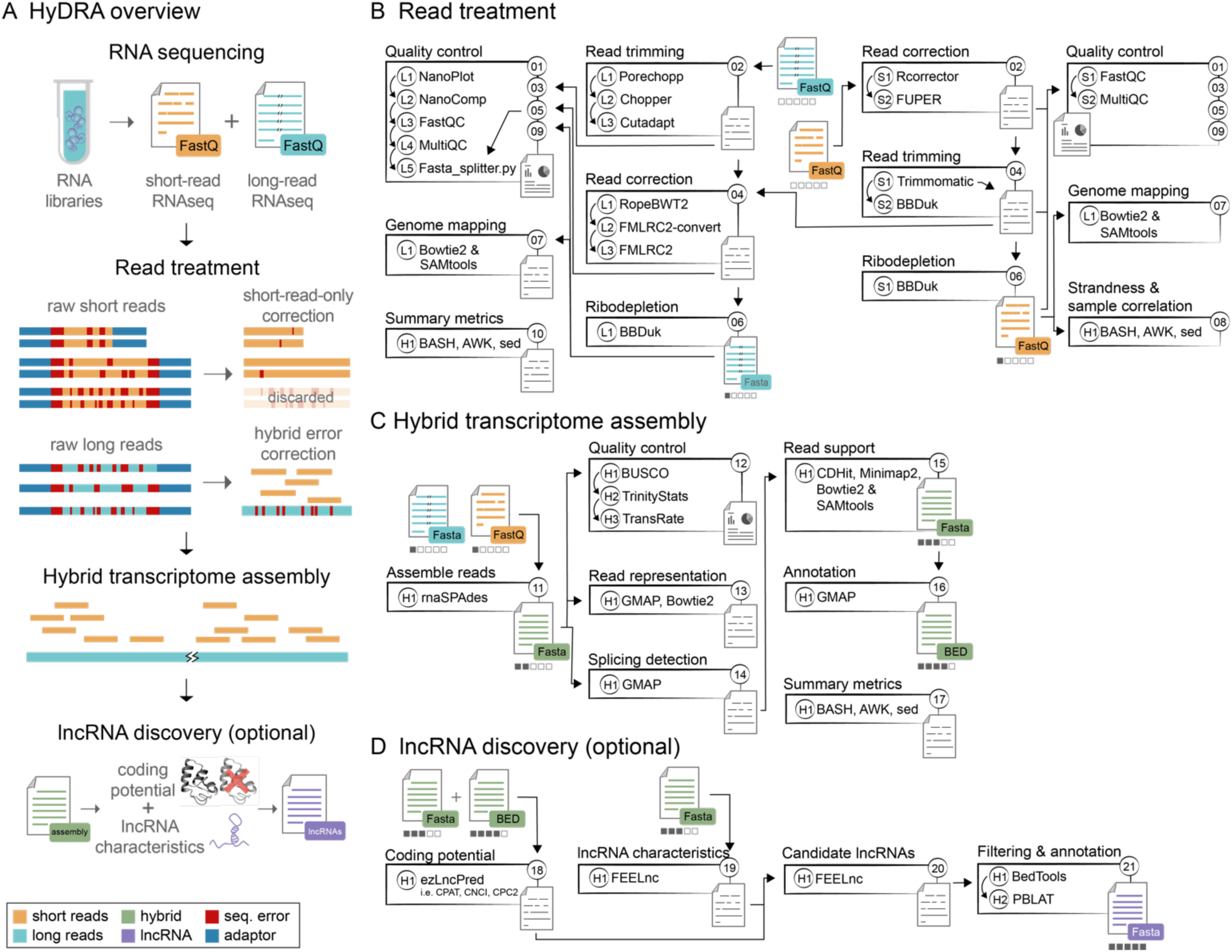
HyDRA. **(A)** Overview of the pipeline, from RNA library preparation to sequencing and availability of raw fastQ files for both short- and long-read samples. **(B)** Both short and long reads first undergo extensive quality control and processing, including hybrid error-correction of long reads and short-read-only correction of short reads. These steps are important to assess low-frequency *k*-mers for error correction and to remove contaminants (e.g. adaptors and reads from ribosomal RNAs). Summary metrics for these steps are printed at the end. **(C)** Treated reads undergo a hybrid *de novo* transcriptome assembly and further filtering and quality assessment. Summary metrics for these steps are printed at the end. **(D)** Optional steps can be performed for the discovery of high-confidence lncRNAs.

**Table 1.**
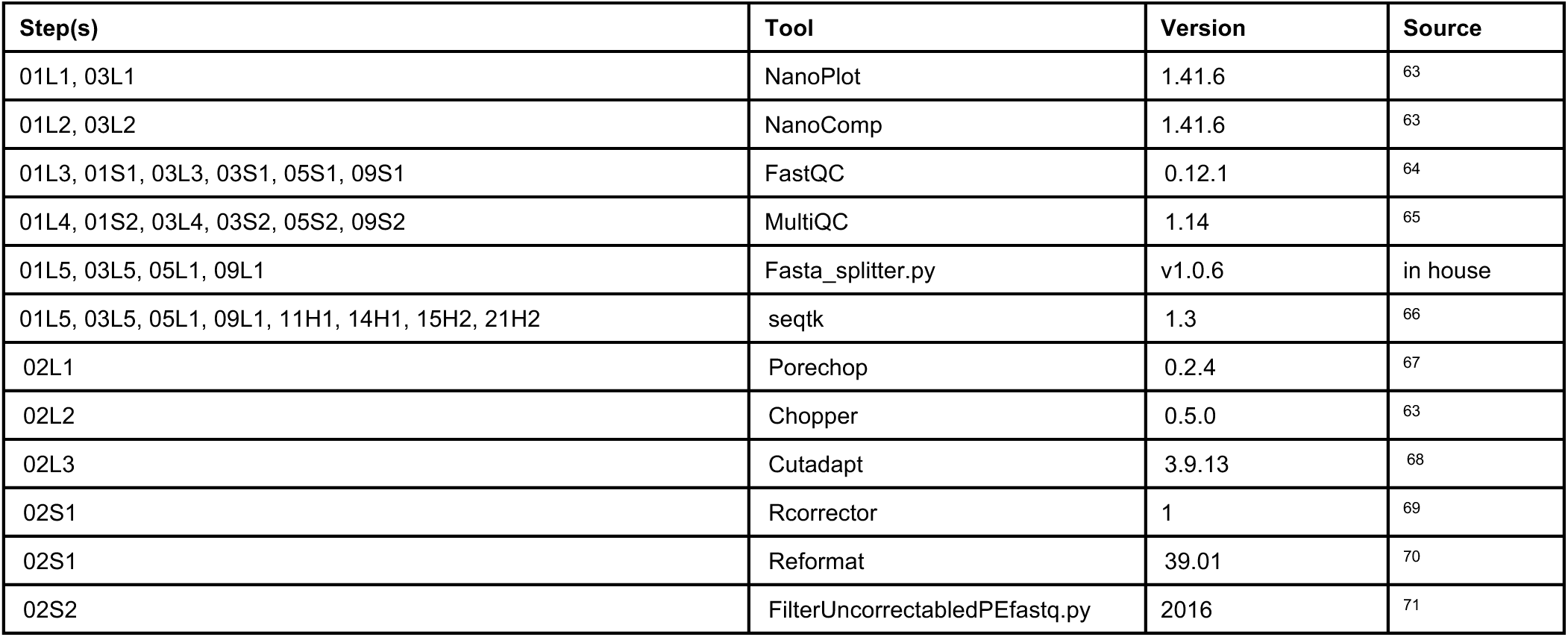
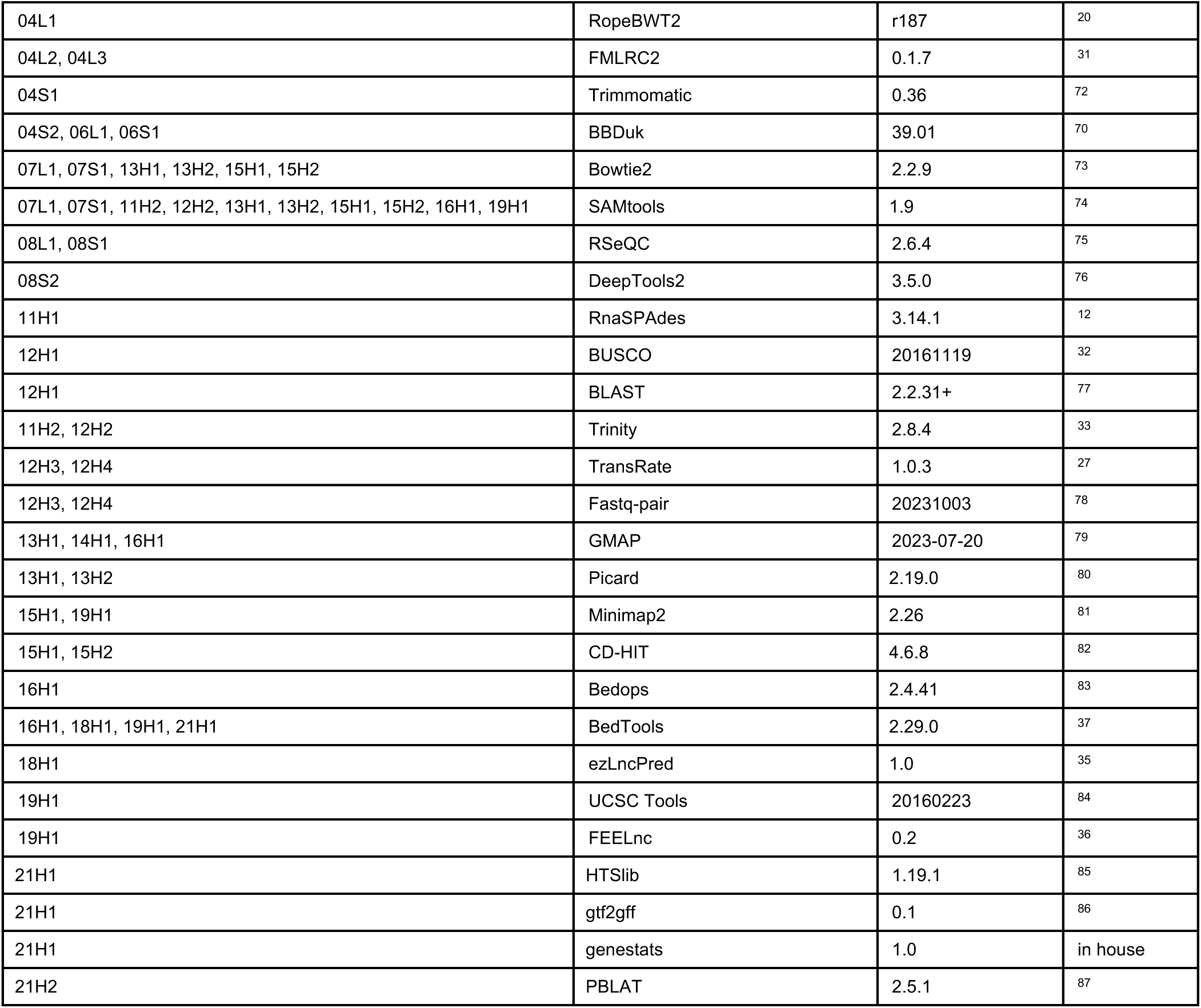
Open-source tools used in HyDRA. Steps are arranged in subroutines specific for long reads (L), short reads (S) and hybrid (H).

#### Read treatment

Sequencing errors are known to introduce artificial nodes in de Bruijn graphs during *de novo* isoform resolution^26^ and interfere with all downstream steps^27,28^. In addition to correcting sequenced reads, common read processing practice includes removing adaptor sequences identified during the raw quality assessment of the data^29^. Traditional tools optimized for short-read data fail to correctly treat longer sequences^7,9,30^. Therefore, HyDRA includes scripts and subroutines carried out by best-performing tools specifically designed to process these data separately (Figure 1B). As a result, HyDRA’s read treatment phase includes 38 scripts that perform the first ten steps. The short-read processing steps of HyDRA follow our previously published *de novo* assembly pipeline, which is currently best-in-class^4^. Processed in parallel, long-read treatment steps are dependent on the pre-processed short-reads for hybrid error correction using FMLRC2 v.0.1.7^10,31^. QC routines are interspersed throughout the read treatment steps to guarantee high read quality, including an in-house Python script (fasta_splitter.py) to assess long-read length, allowing the user to implement personalized cut-offs for ultra-long reads (e.g. > 35 kb). These QC steps were designed to enhance the quality of input read data and are performed after each key processing step.

#### *De novo* assembly

We selected RnaSPAdes v.3.14.1^12^ as the assembler for HyDRA, as it was specifically designed for integrating short and long RNAseq reads and is the only available assembler that uses a genome-independent process (Additional file 2: Table S1, Figure 1C). RnaSPAdes was developed from the foundational algorithms SPAdes and hybridSPAdes, enabling the integration of both paired-end short-read RNAseq data and single-end long-reads, from either PacBio or ONT. This approach facilitates the construction of a high-quality transcriptome assembly that represents full-length transcripts and their alternative isoforms^12^. Next in the HyDRA pipeline, a step is included to remove highly redundant transcripts and differentiate between multiexonic and monoexonic sequences in the assembly. This allows users to set appropriate read support thresholds for each subset of transcripts, with monoexonic transcripts requiring higher read support to differentiate from sequencing noise or genomic DNA contamination. HyDRA uses reads per kilobase per million (RPKM) values from independent short- and long-read alignments to estimate read support, with user-defined thresholds for filtering.

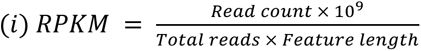

Assembled transcripts are then aligned to the reference transcriptome to identify unannotated transcripts in the custom transcriptome. Similar to our QC routine for input reads, we use a series of biologically supported quality evaluation tools (BUSCO v.20161119^32^, Trinity Stats v.2.8.4^33^ and TransRate v.1.0.3^27^), to assess completeness and other metrics that characterize the generated custom transcriptome. With that, this section of the pipeline includes 9 scripts performing 7 steps, with 3 additional scripts included for short-read-only assembly and processing.

#### LncRNA discovery (optional)

A custom transcriptome assembly can help in the discovery of a variety of transcript types, with lncRNAs representing a substantial portion of the unannotated transcriptome. LncRNAs are highly specific to different tissues, cell types and developmental stages^4,34^. Despite their significance, lncRNAs are often underrepresented in transcriptional studies due to their lack of annotation in reference transcriptomes. This is partially due to short-read RNAseq inherent biases and inability to capture full-length transcripts. Using a combination of long and short reads, HyDRA is well-equipped to facilitate the annotation of lncRNAs. We have therefore included 4 optional steps after the core assembly that allow HyDRA to perform lncRNA discovery. Using a combination of three machine learning models from ezLncPred v.1.0^35^, i.e. CPAT, CNCI, CPC2, we first predict the coding potential of the transcripts identified in the assembly. These transcripts are then assessed in parallel by FEELnc v.0.2 for lncRNA characteristics^36^. FEELnc is a suite of machine learning algorithms that requires the user to supply both a reference annotation of protein-coding transcripts and previously annotated lncRNA transcripts (e.g. GENCODE annotation) to train the model to classify transcripts assembled by HyDRA. A transcript is then considered to be a candidate lncRNA based on FEELnc’s prediction in combination with the predicted absence of coding potential detected by at least two different ezLncPred tools. To remove false-positive lncRNAs (i.e. transcripts that match annotated protein-coding transcripts), HyDRA maps the candidate lncRNAs to the reference transcriptome. Candidate lncRNAs matching protein-coding transcripts with at least 75% identity and a minimum bidirectional overlap of 85% on either strand, are identified as false-positives and removed. Coordinates of candidate lncRNAs are also intersected with protein-coding genes using BedTools^37^, resulting in a final set of high-confidence lncRNAs. Finally, HyDRA maps the candidate lncRNAs to a comprehensive database of confirmed lncRNAs that can be the internal default (containing 112,439 lncRNAs from multiple sources, as described in Bitar *et al.* 2023^4^) or a user-defined database. This allows the user to pinpoint which lncRNAs have been detected for the first time in the custom assembly, and which were already known, either from the reference transcriptome or from additional databases (Additional file 1).

### HyDRA improves the quality of both short- and long-sequenced reads

HyDRA was developed and tested using data obtained from short- and long-read RNAseq on primary and immortalized fallopian tube secretory epithelial cells (FTSEC) and ovarian surface epithelial cells (OSEC) (Additional file 2: Table S2). QC of raw RNAseq data confirmed an expected high median Phred quality of 35.65 for the short-read data, and a median quality of 12.90 for the long-read data (Additional file 2: Table S3; Additional file 3: Figures S2-S3). We used the Illumina NovaSeq™ 6000 for short read sequencing, which has the lowest error rates for high-throughput sequencing^38^. For long-read sequencing, we used ONT with current error rates predicted at 5-10%^9,39^. Common practice with long read RNA sequencing includes the removal of reads below a mean minimum Phred quality of 7 (Q7), which is a much lower threshold than for short-read data. In our dataset, all long-reads were over Q7, with 93% of them surpassing Q10 (Additional file 2: Table S3).

The short-read treatment steps begin with correction of raw reads and subsequent trimming. The majority of the short reads survived the correction step (average of 9.16% uncorrectable reads; Figure 2; Additional file 2: Table S3; Additional file 4) and about 60% of the sequence ends were above the minimum quality set for trimming Q30 (Additional file 2: Table S3), indicating a high quality of corrected and trimmed short reads. Median short-read quality measured in the Phred scale increased from 35.65 to 36.12 after treatment steps were performed (Additional file 3: Figure S1-S2; Additional file 2: Table S3), with a concomitant decrease in the calculated error rate from 1 base in ~3,000 to 1 in ~4,000. In HyDRA, due to the prerequisites of the selected tools, the long-read treatment steps follow the opposite order, with trimming (adaptor removal followed by quality trimming) performed before correction. Median long-read quality measured in the Phred scale showed an increase from 12.90 to 14.50 after trimming. Approximately 5% of the reads were discarded during adaptor removal and quality trimming (Figure 2). From the remaining reads, 99% were above Q10 and 81% were above Q12 (Additional file 3: Figure S3; Additional file 2: Table S3). Next, we integrated the pre-processed short- and long-read sequencing data to perform the hybrid error correction. We observed a balanced base composition in the Burrows-Wheeler transform created from all pre-processed short reads. However, we consistently noticed that RopeBWT2^20^, one of the tools used in the hybrid error correction steps (more details on Additional file 1) outputs a base count report in which thymine base counts and N (undefined) base counts are swapped. This has been addressed in HyDRA which now outputs the correct base counts to the user (Additional file 2: Table S3). Despite base quality information being lost after long-read correction, no sequences were discarded at this point, implying that all reads that survived trimming were corrected and kept for further processing (Additional file 2: Table S3).

**Figure 2.**
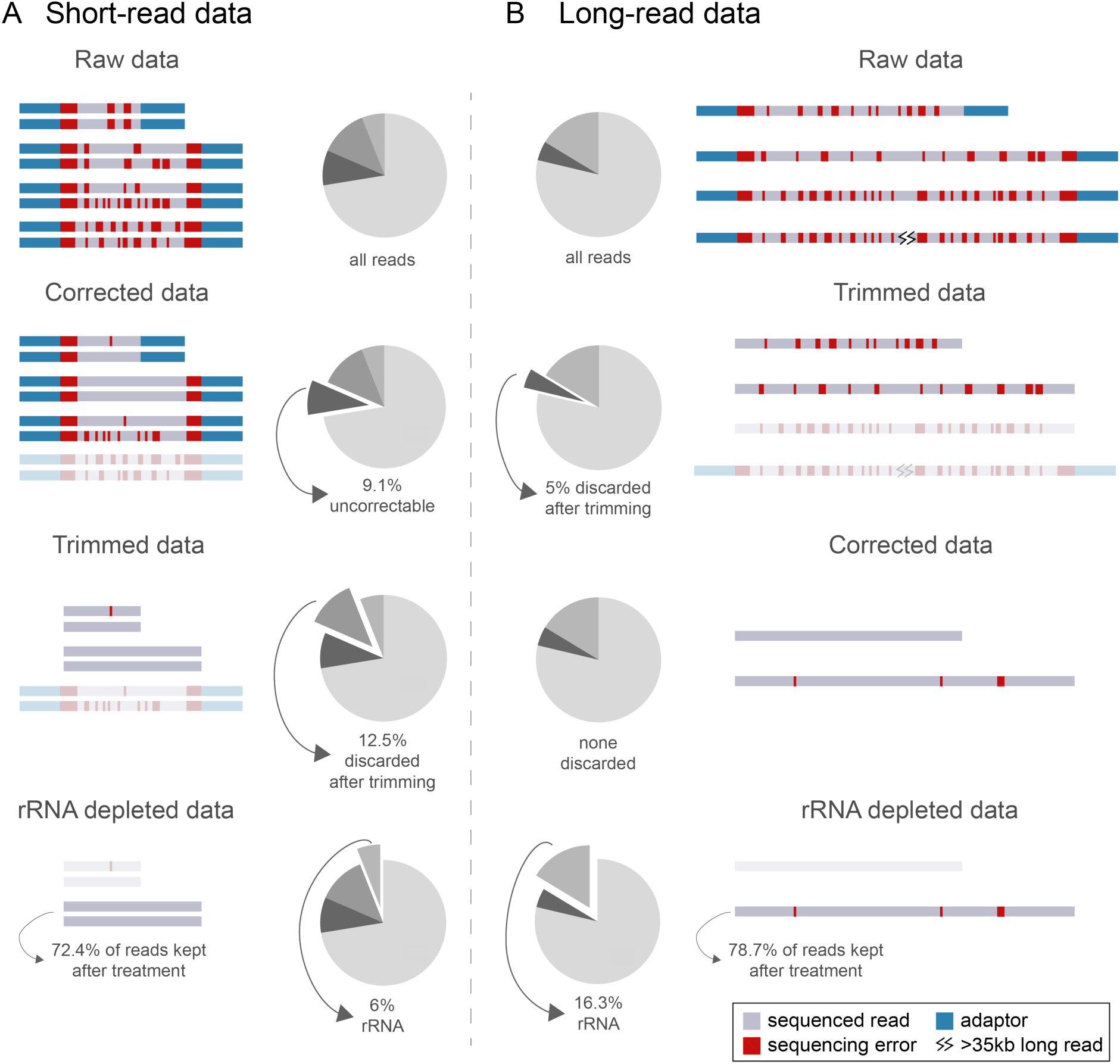
Short- and long-read data treatment in HyDRA. **(A)** Paired-end short-read data, followed by pie charts highlighting the proportion of reads discarded in each step relative to the total number of raw reads. **(B)** Single-end long-read data, preceded by pie charts highlighting the proportion of reads discarded in each step relative to the total number of raw reads.

Most RNAseq library preparation protocols include a ribodepletion or poly(A) selection step, but ribosomal RNAs (rRNAs) still represent a large portion of the sequenced data^29^. These are considered cognate contaminants, meaning they are reads originating from undesired RNA types and must be removed prior to the *de novo* assembly. Using a database of known rRNAs sequences (Additional file 2: Table S1), we have included a step in HyDRA where pre-processed reads are computationally filtered to remove ribosomal contamination. During quality assessment with FastQC, two long-read sequences identified as overrepresented were confirmed through BLAT searches to be human rRNAs^40^. On average, short-read data contained 6.00% of rRNA-derived reads and long-read data contained 16.30% (Figure 2; Additional file 2: Table S3). These numbers align with expected rRNA sequencing levels, even after ribodepletion during library preparation^41^.

### Long-read sequences of up to 35 kb were kept and used for assembly

The longest known human transcript is *TTN* (titin), with 109,224 nt^42^, thus we anticipated that certain long-read sequences in our dataset would be significantly longer than the reported average of 1-3 kb^18, 19^, potentially including ultra-long reads over 100 kb in length. Indeed, although the average length of the raw long reads was 1024 nt, the longest was 103,744 nt (Additional file 3: Figure S4A-C; Additional file 2: Table S3). FastQC analysis indicated that, while reads with up to ~30,000 nt had the expected base composition (i.e. balanced proportions of A, T, C and G), longer sequences presented a distinctively biased pattern of nucleotide composition, rich in thymines and guanines (Additional file 3: Figure S4B). In light of this, we implemented an additional QC routine to analyze sequence lengths throughout the pipeline and allow users to remove sequences with unexpected nucleotide composition (fasta_splitter.py; Additional file 3: Figure S4D). After all read treatment steps were performed, the eight remaining longest sequenced reads (Additional file 2: Table S3; 30-35 kb), were aligned to the GRCh38 reference genome using BLAT^40^ (Figure 3). All were confirmed as valid human sequences, aligning to *AHNAK* (desmoyokin), *DST* (dystonin) or *LYST* (lysosomal trafficking regulator). All treated long-read sequences were used for *de novo* assembly, including those reaching 35 kb. Importantly, this maximum length is dependent on the input data, the tissue(s), developmental stage and species being analyzed. Additionally, through fasta_splitter.py, HyDRA gives users the option to strictly keep sequences that have up to *n* nt in length.

**Figure 3.**
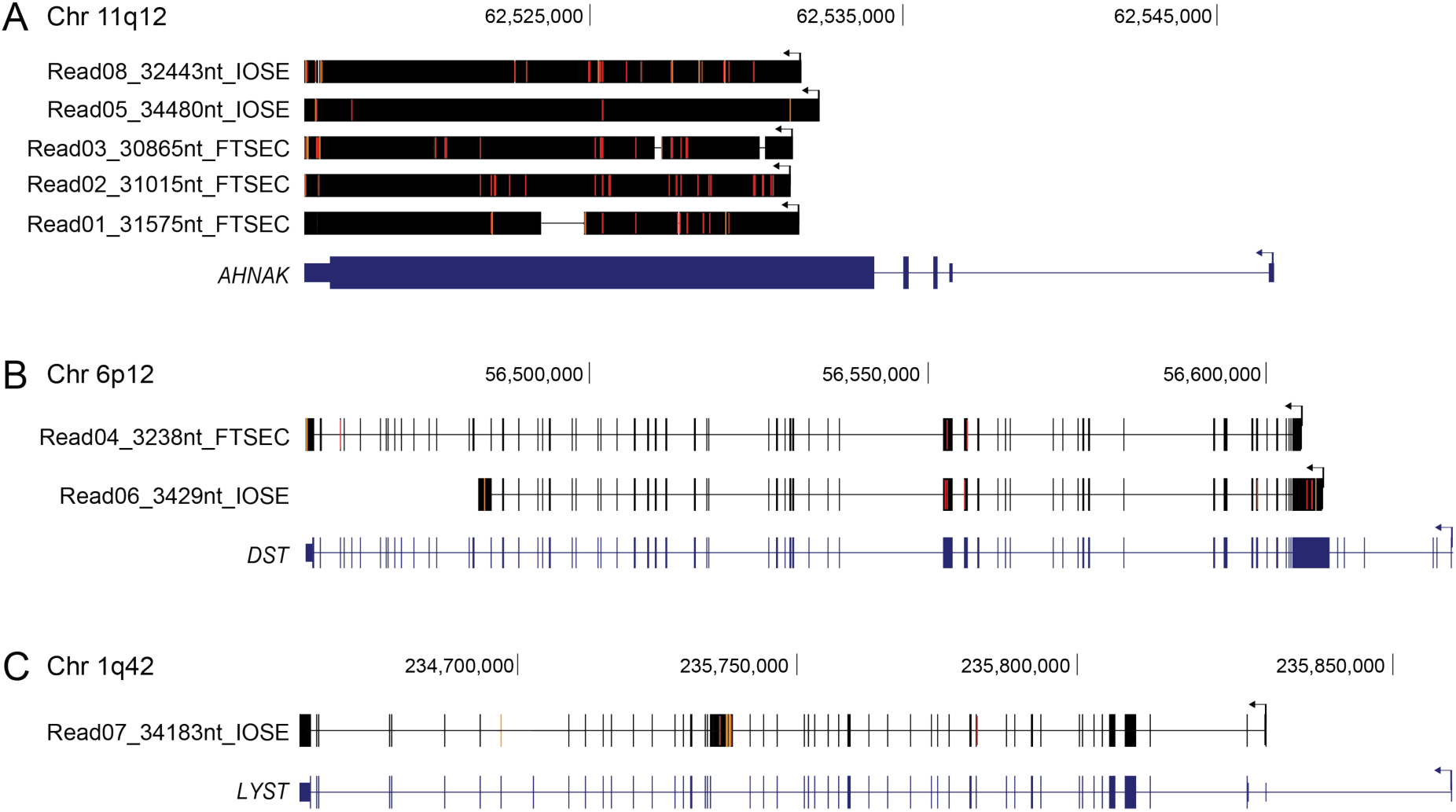
Treated long-read sequences reaching up to 35 kb aligned to the human genome using BLAT. These eight pre-processed reads (four from FTSEC and four from OSEC samples) were aligned against the human genome (GRCh38) in a UCSC BLAT search to confirm they were valid sequences. The sequencing reads that aligned to **(A)** *AHNAK*, **(B)** *DST* or **(C)** *LYST*.

### Hybrid transcriptome assembly performs better than short-read-only approaches

To assess HyDRA’s assembly, a short-read-only assembly was created by combining the treated short reads as an input for Trinity v.2.8.4^33^ with normalized read coverage at 50 to prevent fragmented transcripts^4^. This assembly was subjected to the same processing steps as the hybrid counterpart. To evaluate the quality of both assemblies, we used a subset of the metrics reported in a recent benchmark study and respective normalized score (0-1)^43^. These metrics included transcript length, N50, reference coverage, open reading frame (ORF) percentage, undefined base count and conserved orthologs representation. Based on the normalized score, the HyDRA generated assembly performed 31% better than the best performing short-read-only approach^4^, and outperformed the top-ranked *de novo* assembly tool alone by 41% (Figure 4A; Additional file 2: Table S4)^43^. Our hybrid approach generated 857,736 transcript sequences, reaching up to 67,466 nt, with an average transcript length of 2,409 nt and GC content of ~44% (Additional file 2: Table S4), which aligns with the reported human GC content of coding (~52%) and noncoding (~44%) isoforms^44^.

**Figure 4.**
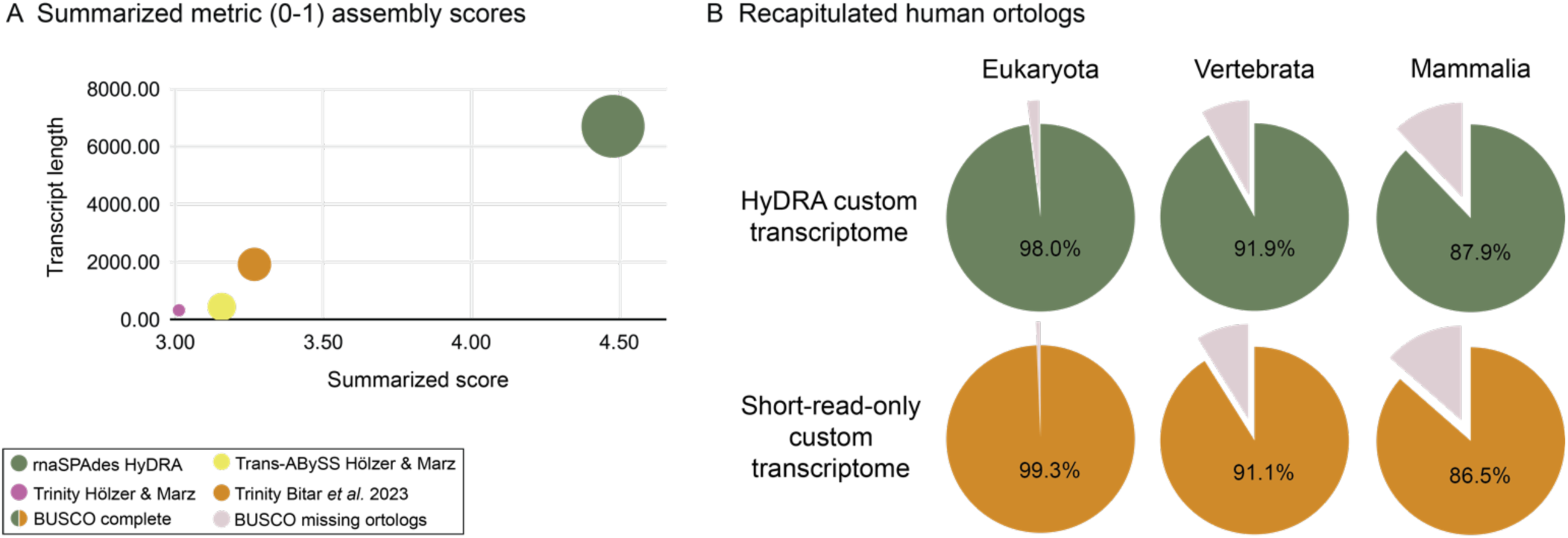
Overall assessment of the HyDRA-generated assembly. **(A)** Normalized assembly scores. Bubbles sizes vary according to N50 value. The graph shows scores for the assemblies produced by i) the best (Trans-ABySS) and ii) second best *de novo* assembly tools alone (Trinity)^43^; iii) Bitar *et al.* 2023^4^ pipeline; iv) HyDRA. Both (i) and (ii) were based on data described by Hölzer and Marz’s^43^ and (iii) and (iv) based on data described here (from human ovarian and fallopian tube samples). **(B)** HyDRA’s assembly completeness from BUSCO analysis.

In terms of assembly contiguity, N50 is an important metric defined as the length of the sequence at which 50% of the total assembly size is contained in sequences of at least that length. HyDRA produced an assembly with N50 of 6708 nt, which reflects how a hybrid approach can represent full-length human transcripts (Additional file 2: Table S4). For comparison, Hölzer and Marz’s best performing assembler produced a transcriptome with an N50 of 441 nt (15.21 times smaller than HyDRA’s assembly)^43^ and Bitar *et al.* 2023 an N50 of 1383 (4.85 times smaller than HyDRA’s assembly)^4^. The highest N50 observed by Hölzer and Marz^43^ was 2381 nt (2.82 times smaller than HyDRA’s assembly), but this study showed that the assembler performed poorly compared to the other tools and metrics. To investigate the contribution of adding long reads to transcriptome assembly, we used the Bitar *et al.* 2023^4^ pipeline to create an assembly based only on our short-read data, comparing it with HyDRA’s hybrid assembly. The calculated N50 of the short-read-only assembly was three times smaller than HyDRA’s and the assembly had double the number of transcripts. This suggests HyDRA can generate less fragmented assemblies, that are likely to better recapitulate full-length transcripts, while maintaining high overall quality. Similar to the N50, the N90 metric corresponds to the transcript length at which 90% of the total assembly size is contained in sequences of at least that length. Using our hybrid approach, we achieved an N90 of ~1000 nt, meaning that 90% of the transcripts in the HyDRA assembly are sequences matching the average human transcript length^17^. This demonstrates the overall contiguity of the produced custom transcriptome (Additional file 3: Table S4).

The HyDRA-generated assembly accurately recapitulated several aspects of the human transcriptome. For example, BUSCO analysis revealed > 98% of the eukaryotic (297/303), 91.9% of the vertebrata (2376/2586) and 87.8% of the mammalian (3606/4104) conserved orthologs were captured in our hybrid assembly, indicating overall completeness (Figure 4B). These values were similar to those obtained from the short-read-only assembly (Additional file 2: Table S4). According to TransRate, the custom hybrid assembly covered 24% of the reference human transcriptome (GENCODE), which is comparable to the 23-26% observed in the best performing assembler tools found by Hölzer and Marz^43^ (Additional file 2: Table S4). For perspective, HyDRA’s transcripts cover approximately 12% of the reference genome while the exons and UTRs in the reference annotation (GENCODE v36) cover approximately 5% (genome coverages were calculated with BedTools genomecov).

### Splicing assessment showed 30% of transcripts to be multiexonic

Most assemblies to date disregard monoexonic transcripts, but recent evidence has shown this class contains conserved lncRNAs of functional relevance^45,46,47,48^. Similar to Bitar *et al.* 2023^4^, we have kept the monoexonic transcripts in our assembly, as long as they had high read support. Transcripts aligned to the reference human genome (GRCh38) were classified as monoexonic or multiexonic according to the presence of ‘N’ tags in the alignment file. A minimum length of 50 nt was defined to differentiate introns from insertions and deletions (indels), which aligns with current knowledge about human introns. Before filtering out low read support transcripts, our hybrid assembly showed a ratio of multiexonic:monoexonic transcripts of 3:7 (~232,000 transcripts were classified as multiexonic and ~619,000 as monoexonic; Figure 4C; Additional file 2: Table S4). For comparison, the short-read-only approach showed a ratio of 3:17 (~143,000 were multiexonic and ~740,000 were monoexonic), likely reflecting the power of long reads to resolve transcript architecture and improve overall isoform assembly.

Removing transcripts with low read support helps remove technical artifacts and transcriptional leakage products, as well as problematic transcripts arising from misassembly. As HyDRA integrates short and long reads, read support for each transcript was calculated based on a combination of both subsets, which is computationally and biologically challenging. In HyDRA, redundant transcripts are collapsed prior to read support calculations. This redundancy reduction step removed ~8,500 transcripts from the original assembly (Additional file 2: Table S4). From the remaining ~224,500 multiexonic and ~618,000 monoexonic transcripts, ~189,000 (84.34%) and ~13,000 (2.12%) respectively, passed the more permissive RPKM cut-off for read support (0.3 and 3 RPKM). As expected, the number of supported transcripts was much lower when using the stricter RPKM cut-off (1 and 5 RPKM), with a 90% decrease in multiexonic (~20,000) and 50% decrease in monoexonic transcripts (~7,300) passing the filtering step. Since we previously validated transcripts with low read support by qPCR, confirming that the less stringent cut-off still identifies bona fide transcripts^4^, we opted to use these transcripts for further analysis. The ratio of multiexonic to monoexonic transcripts in the assembly is 1:14, maintaining the expected lncRNA ratio observed in the Telomere-To-Telomere (T2T) human genome (T2T-CHM13). In total, the final filtered custom assembly consists of 202,459 transcripts, providing a comprehensive representation of the normal ovarian and fallopian tube transcriptome.

### Identification of unannotated lncRNAs in HyDRA’s custom transcriptome

To assess the coding potential of the 202,459 transcripts, we ran three machine learning models from the ezLNCpred package, CPAT, CNCI and CPC2^35^. On average, at least two of the models agree on 61% of the noncoding predictions (93,899), suggesting that these methods are more effective at confirming the absence of ORFs rather than detecting their presence. Additionally, 26.6% (40,969) had no coding potential detected by any of the three models (Figure 5A; Additional file 2: Table S5). A total of 47,281 transcripts were predicted by at least two models as noncoding and not by any model as protein-coding. We decided to include all 93,899 transcripts predicted as noncoding by at least two of the tools in our further analysis (Figure 5B; Additional file 2: Table S5).

**Figure 5.**
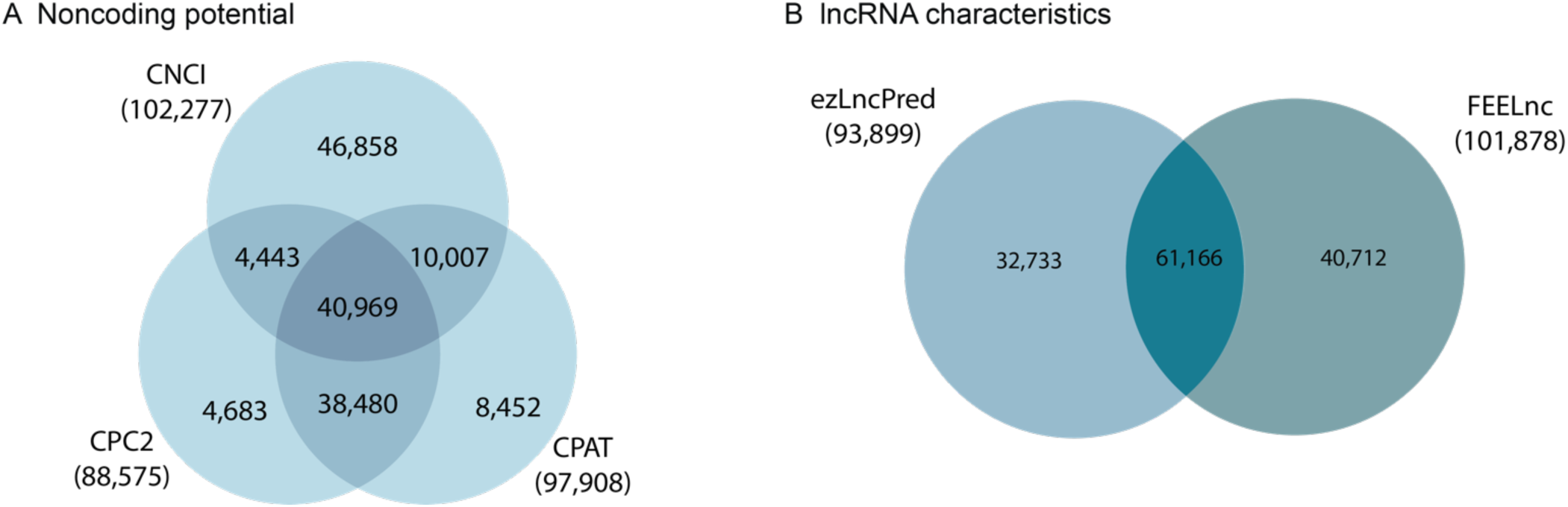
LncRNA discovery. **(A)** Intersection of lncRNA candidates predicted by three different ezLncPred machine learning models (CNCI, CPAT and CPC2). **(B)** Intersection between FEELnc lncRNA predictions and the list of candidates predicted by at least two of the ezLncPred machine learning models.

FEELnc was trained using the GENCODE GRCh38 transcriptome annotation of protein-coding and lncRNA transcripts^49^. This enabled us to identify which of the 202,459 transcripts assembled by HyDRA had lncRNAs characteristics. To ensure robust predictive capability, we exclusively used experimentally validated GENCODE lncRNAs. FEELnc predicted 101,878 transcripts to be lncRNAs (Additional file 2: Table S5). This represents about half of the total number of transcripts in our custom transcriptome, indicating that lncRNAs constitute a significant proportion of expressed transcripts in normal ovarian and fallopian tube tissues. Importantly, more than 60% of these (61,166) lncRNAs were supported by at least two of the three ezLncPred machine-learning models (Figure 5B).

The majority of these candidate lncRNAs were multiexonic, which may reflect the biased training dataset. From the 61,166 candidate lncRNAs, 629 showed a bidirectional overlap of at least 85% with a protein-coding transcript and a minimum identity of 75%. We believe these to be either false-positives that our methods failed to detect, or noncoding isoforms of protein-coding genes. Although monoexonic and sense genic transcripts (i.e. those overlapping protein-coding genes in the same strand) are functionally relevant, it is difficult to differentiate these from technical artifacts or transcriptional leakage. Furthermore, sense genic lncRNAs cannot easily be uncoupled from the corresponding protein-coding gene, and we have included a restrictive alignment step to facilitate their removal. From the remaining 60,537 lncRNAs, 228 were already annotated in GENCODE (GRCh38: 19 multiexonic; 1 monoexonic) or known to the database of over 112,000 lncRNA transcripts (124 multiexonic; 84 monoexonic). We also intersected the coordinates of the remaining 60,309 lncRNAs with those of protein-coding genes using BedTools^37^. This allowed us to further remove 7,615 exon-overlapping transcripts, resulting in the identification of 53,551 high-confidence lncRNA transcripts. Importantly, HyDRA was designed to split monoexonic and multiexonic sequences after assembly, which allows users to easily focus on either or both sets of transcripts.

To demonstrate the functional relevance of these 53,551 high-confidence lncRNA transcripts from the normal ovaries and fallopian tubes, we assessed their expression profiles in a different subset of sequenced RNA samples (unrelated but biologically similar to those used for assembly). We used publicly available RNAseq data from normal and cancerous ovarian and fallopian tube tissues found in the RNA Atlas project^50^. This revealed that 27,257 (44.56%) lncRNAs were expressed in at least one of the samples, demonstrating HyDRA’s efficient assembly of both annotated and unannotated lncRNA transcripts, through the integration of long- and short-read RNAseq data.

## CONCLUSIONS

Here we present HyDRA, a comprehensive pipeline that integrates short- and long-read sequencing data for a true-hybrid *de novo* transcriptome assembly and lncRNA discovery. We used deep, short- and long-read RNAseq from ovarian and fallopian tube epithelial cells samples to develop, validate and assess the efficacy of the pipeline in generating a high-quality custom transcriptome. We have shown that HyDRA’s assembly performed > 40% better than the top-ranked stand-alone *de novo* transcriptome assembly tool and > 30% better than our recent best-in-class short-read-only approach^4^. Based on this custom assembly, we identified 61,166 candidate lncRNAs, among which 60,309 have not been previously annotated and 53,551 showed no overlap with protein-coding transcripts. In summary, HyDRA is a high-performing hybrid-assembly tool capable of facilitating accurate transcriptome reconstruction and advancing lncRNA annotation.

## MATERIALS AND METHODS

### RNAseq sample preparation and sequencing

RNA was extracted from primary and immortalized fallopian tube secretory epithelial cells (FTSEC) and ovarian surface epithelial cells (OSEC) using the QIAGEN RNeasy Plus Mini kit (Additional file 2: Table S2). One microgram of total RNA was rRNA depleted with the Ribo-Zero™ Plus kit according to the manufacturers’ instructions (Illumina). Short-read RNAseq libraries were prepared using the Truseq Stranded mRNA Library Prep Kit (Illumina), and sequenced at high depth (PE150, *>* 75 million reads per sample; Additional file 2: Table S2) on the Illumina Novaseq™ 6000 (Australian Genome Research Facility, Melbourne, Australia). High sequence depth is considered best practice for lncRNA discovery, as they are often expressed at low levels and have poor isoform representation^51^. For long-read sequencing, cDNA was extracted from one FTSEC and one OSEC cell line (Additional file 2: Table S2). ONT cDNA libraries were generated with polyadenylation enrichment and SQK-NBD114.24 native barcoding kit at the Garvan Institute’s Nanopore Sequencing Facility (Australia). Samples were barcoded using the supplied PCR barcodes and sequenced at high depth (*>* 69 million reads per sample; Additional file 2: Table S2) on the PromethION™ P48 flowcells (FLO-PRO114M - R10.4.1). The slow5 files were base-called using Guppy v.6.4.6+ae70e8f and MinKNOW v.22.12.5 by the Garvan Institute’s Nanopore Sequencing Facility (Australia).

### Databases and reference genome versions

We used the human genome GCRh38 release 79^49^ for lncRNA identification and annotation), and the T2T-CHM13 genome^52^, for long-read effects on the produced transcriptome assembly. The GENCODE annotation for GCRh38 was used as the reference transcriptome. A previously published database of 169 human rRNA sequences^4^, with the addition of two sequences identified from our FastQC analysis (Additional file 2: Table S2), was used to filter pre-processed reads for ribosomal contamination. To identify which of the discovered lncRNAs were known and which were novel, we used all annotated lncRNAs in GENCODE GRCh38 together with a comprehensive database of > 112,000 known lncRNA genes from 8 public repositories (BIGtranscriptome, MiTranscriptome and LNCipedia from lncRNAKB^53^; Cabili *et al.* 2011^54^; CancerSEA^55^; Lanzos *et al.* 2017^56^; LncRNADisease^57^ and RNAcentral^58^), as described in Bitar *et al.*^4^. Experimentally confirmed protein-coding and lncRNAs annotated in the same GENCODE version were used to train the machine-learning algorithms FEELncfilter, FEELnc_codpot and FEELnc_classifier^36^.

### Parameters used for the ovarian and fallopian tube custom assembly

A comprehensive list of the parameters used in each step is available in Additional file 2: Table S1). Importantly, we defined both a restrictive and permissive set of cut-offs for read support. A strict read support of 3 RPKM was enforced for monoexonic transcripts, but we relaxed the cut-off to 0.3 RPKM for multiexonic transcripts. This is in agreement with^4^ and maintained the expected ratio of multiexonic to monoexonic lncRNAs of 14:1, consistent with annotations based on the T2T genome. A stricter threshold of 5 RPKM and 1 RPKM, respectively, was also tested. Importantly, lncRNAs expressed at ~0.5 RPKM had previously been experimentally confirmed by our group with an 80% success rate^59^.

### Expression profiles of lncRNA transcripts

RNAseq analysis was run based on the GRADE (General RNAseq Analysis for Differential Expression) pipeline^4^. Modifications to these scripts now allow the user to quantify reads based on any user-provided transcriptome sequence and are available at^60^. The expression profiles of lncRNAs were assessed in (i) ovarian and fallopian tube whole tissue; (ii) high-grade serous ovarian carcinoma (HGSOC) tumor samples, including homologous recombination (HR)-deficient and HR-proficient; and (iii) HGSOC cell lines. Public RNAseq of ovarian and fallopian tube samples, sequenced at high depth, were obtained from the RNA Atlas project^50^.

### Bioinformatics tools used for HyDRA development

Our pipeline integrates the currently available open-source tools in BASH scripts using basic UNIX commands to write and submit portable batch system (PBS) jobs. Our scripts were designed to run in a high-performance computer (HPC) where computational tasks are allocated in a PBS. However, general command lines are also available for users that wish to run the pipeline on a different system (See Availability of Data and Materials). Resources used for pipeline development are indicated in each script and can be controlled by the user according to their available computational power. The length evaluation script, fasta_splitter.py, was developed in Python 2.7+. To assess HyDRA’s assembly, a short-read-only assembly was created by combining the treated short reads as an input for Trinity 2.8.4^33^ with normalized read coverage at 50 to prevent fragmented transcripts^4^.

HyDRA was developed from 39 open-source tools and runs through BASH scripts (Table 1). A comprehensive list of the all tools used in each step is available in Additional file 2: Table S1). BLAT (BLAST-like alignment tool implemented at the UCSC genome browser) searches^40^ were performed to confirm rRNA sequences, long-read sequences and investigate identified lncRNAs. Plots were produced either directly by the underlying tools (referenced in text), with Python 2.7+ script available at HyDRA GitHub repository^61^). Venn Diagrams were generated with InteractVenn^62^. Figures were edited in Adobe Illustrator v.28.5.

## SUPPLEMENTARY INFORMATION

Additional file 1: Complete HyDRA pipeline description

Additional file 2: Tables S1-S5

Additional file 3: Figures S1-S4

Additional file 4: Pipeline validation

## AUTHORS CONTRIBUTIONS

I.A. and M.B. designed and directed the study and interpreted the data. I.A. developed the pipeline, performed all dry-lab analyses, critically reviewed the computational code, created and maintains the GitHub repository. X.L. prepared the RNA for short- and long-read RNA sequencing. I.A., J.D.F., S.L.E., and M.B. conceived the project and wrote the manuscript with contributions from all authors.

## FUNDING

This work was supported by grants from the National Health and Medical Research Council of Australia (NHMRC; 2019101) and The Donald and Joan Wilson Foundation. I.A. was supported by a QIMR Berghofer International PhD scholarship and a University of Queensland (UQ) Research Training scholarship. J.D.F. was supported by an NHMRC Investigator Grant (2016826).

## AVAILABILITY OF DATA AND MATERIALS

The HyDRA pipeline in source and executable form is available without charge for nonprofit, academic, and personal uses under an MIT license at the HyDRA GitHub repository HyDRA^61^ Zenodo. HyDRA was developed using a x86_64 GNU/Linux operational system in a HPC environment. Expression profiles of lncRNAs discovered from the hybrid custom transcriptome assembly produced were obtained with GRADE2 (General RNAseq Analysis for Differential Expression, version 2), an open-source pipeline under an MIT license and publicly available on GitHub^60^ and Zenodo. Raw short and long RNAseq data generated in this study have been deposited in the National Center for Biotechnology Information’s Gene Expression Omnibus (GEO) and are accessible through GEO series accession number XXX000000. Previously published total RNAseq data from the RNA Atlas project under GEO series accession number GSE138734^50^ has been used to perform RNAseq analysis (SRR accession codes are available at Additional file 2: Table S2).

## COMPETING INTERESTS

The authors declare that they have no competing interests.

## Notes

### Competing Interest Statement

The authors have declared no competing interest.

